# An estimate of the longitudinal pace of aging from a single brain scan predicts dementia conversion, morbidity, and mortality

**DOI:** 10.1101/2024.08.19.608305

**Authors:** Ethan T. Whitman, Maxwell L. Elliott, Annchen R. Knodt, Wickliffe C. Abraham, Tim J. Anderson, Nick Cutfield, Sean Hogan, David Ireland, Tracy R. Melzer, Sandhya Ramrakha, Karen Sugden, Reremoana Theodore, Benjamin S. Williams, Avshalom Caspi, Terrie E. Moffitt, Ahmad R. Hariri, The Alzheimer’s Disease Neuroimaging Initiative

## Abstract

To understand how aging affects functional decline and increases disease risk, it is necessary to develop accurate and reliable measures of how fast a person is aging. Epigenetic clocks measure aging but require DNA methylation data, which many studies lack. Using data from the Dunedin Study, we introduce an accurate and reliable measure for the rate of longitudinal aging derived from cross-sectional brain MRI: the Dunedin Pace of Aging Calculated from NeuroImaging or DunedinPACNI. Exporting this measure to the Alzheimer’s Disease Neuroimaging Initiative and UK Biobank datasets revealed that faster DunedinPACNI predicted participants’ cognitive impairment, accelerated brain atrophy, and conversion to diagnosed dementia. Underscoring close links between longitudinal aging of the body and brain, faster DunedinPACNI also predicted physical frailty, poor health, future chronic diseases, and mortality in older adults. Furthermore, DunedinPACNI followed an established socioeconomic health gradient with people of lower socioeconomic status showing faster DunedinPACNI. Associations between DunedinPACNI and cognitive impairment were replicated in BrainLat, a sample of Latin American patients with dementia. When compared to brain age gap, an existing MRI aging biomarker, DunedinPACNI was similarly or more strongly related to clinical outcomes. DunedinPACNI is a “next generation” MRI measure that will be made publicly available to the research community to help accelerate aging research and evaluate the effectiveness of dementia prevention and anti-aging strategies.

---

Aging is the gradual, progressive, and correlated decline of multiple organ systems over decades. Longitudinal studies provide evidence for substantial individual variation in the rate of aging; people born in the same year can age slower or faster than their peers^1–3^. Furthermore, aging itself is increasingly regarded as a potentially preventable cause of chronic disease. Accordingly, accurate and reliable measures of how fast a person is aging are needed to effectively study how individual variation in the rate of aging contributes to disease risk and to evaluate interventions intended to slow aging before irreversible decline^4–8^.

Age-sensitive alterations in DNA methylation, referred to as “epigenetic clocks,” are currently the most widely used measures for estimating individual differences in aging^4,9,10^. First-generation epigenetic clocks were trained on chronological age^11,12^, but the more precisely they predicted chronological age, the less well they predicted clinical outcomes^13,14^. In response, second-generation clocks were trained on measures of health that predict mortality in older people^15–17^. However, these clocks were trained on cross-sectional phenotypes in multi-age samples, not on longitudinal observations of the same person as has been recommended in geroscience^5,18^. This limitation led to the development of a third-generation, longitudinal approach to measuring aging.

We previously adopted this longitudinal approach in the Dunedin Study, which has followed a population-representative sample of 1,037 people born in the same year (1972-1973) from birth to age 45^19^. Across two decades (ages 26, 32, 38, and 45 years), we repeatedly measured 19 biomarkers of cardiovascular, metabolic, renal, immune, dental, and pulmonary functioning. By averaging the decline in the trajectories of these biomarkers, we operationalized the theoretical construct of biological aging into a specific measure that we called the Pace of Aging^2^. We subsequently developed an epigenetic clock that accurately and reliably estimates the Pace of Aging: the Dunedin Pace of Aging Calculated from the Epigenome or DunedinPACE^20^. Because DunedinPACE is calculated from a single timepoint measurement of DNA methylation, it has been rapidly adopted by aging studies where it has been associated with signs of accelerated brain aging, morbidity, and mortality^20–25^. However, it has not been possible to export DunedinPACE or other epigenetic clocks to studies lacking DNA methylation data. This includes many neuroimaging studies of brain aging and neurodegenerative diseases such as Alzheimer’s disease.

Current neuroimaging-based approaches to measure aging, akin to first-generation epigenetic clocks, involve training models to predict chronological age from variability in MRI measures of brain structure in large multi-age samples^26–30^. Researchers then typically quantify a “brain age gap,” which reflects the difference between a participant’s predicted and actual chronological age. A positive brain age gap is interpreted as evidence of accelerated brain aging. As with first-generation epigenetic clocks, these age-deviation approaches unavoidably mix model error (e.g., historical differences in environmental exposures, survivor bias, disease effects, measurement bias) with a person’s true rate of biological aging^31–34^.

Here, using a single T1-weighted MRI scan collected at age 45 in the Dunedin Study, we describe the development and validation of a novel brain MRI measure for the Pace of Aging (**Figure 1A-C**). We call this measure the Dunedin **P**ace of **A**ging **C**alculated from **N**euro**I**maging or “DunedinPACNI.” Using data from the Human Connectome Project we evaluated the test-retest reliability of DunedinPACNI. Exporting the measure to the Alzheimer’s Disease Neuroimaging Initiative (ADNI), the UK Biobank, and BrainLat, we conducted a series of preregistered analyses (link: https://rb.gy/b9x4u6) designed to evaluate the utility of DunedinPACNI for predicting multiple aging-related health outcomes (**Figure 1D**). To benchmark our findings, we compared effect sizes for DunedinPACNI to those for brain age gap^35^. DunedinPACNI is the first brain-based measure trained to directly estimate longitudinal aging of non-brain organ systems. If DunedinPACNI does indeed estimate individual differences in the rate of aging, it would add evidence for close links between brain integrity and whole-body aging and establish neuroimaging as a powerful tool for measuring aging; not just of the brain, but of the entire body^36^.

**Figure 1.**
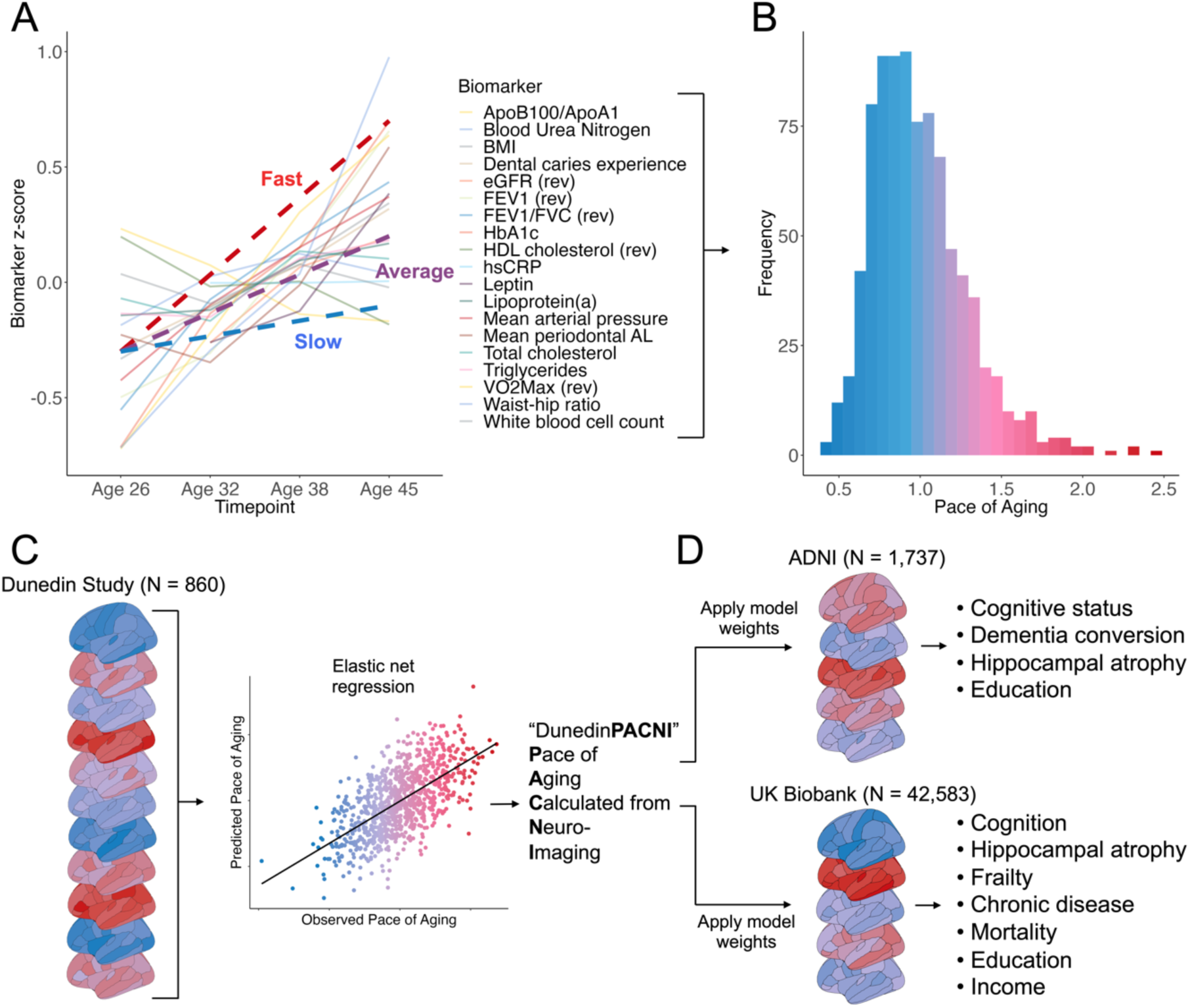
Schematic overview of study methods. **A.** Plot of mean scores for all 19 biomarkers comprising the Pace of Aging across four waves of observation at ages 26, 32, 38, and 45 years in the Dunedin Study. Hypothetical individual trajectories are shown for a person with relatively Slow, Average, and Fast Pace of Aging from ages 26 to 45. **B.** Distribution of Pace of Aging composite scores in Dunedin Study members at age 45. Warmer colors represent a faster Pace of Aging and cooler colors represent a slower Pace of Aging. **C.** A single T1-weighted MRI scan collected from 860 Dunedin Study members at age 45 was used to train an elastic net regression model to predict the Pace of Aging. We call the resulting measure the Dunedin Pace of Aging Calculated from NeuroImaging, or DunedinPACNI. **D.** Regression weights from the DunedinPACNI model developed in the Dunedin Study were applied to T1-weighted MRI scans collected in the Alzheimer’s Disease Neuroimaging Initiative and UK Biobank datasets to derive DunedinPACNI scores. Those scores were then related to aging-related phenotypes. Abbreviations: ADNI = Alzheimer’s Disease Neuroimaging Initiative, AL = attachment loss, Apo = apolipoprotein, BMI = body mass index, eGFR = estimated glomerular filtration rate, FEV = forced expiratory volume, FVC = forced vital capacity, HbA1c = glycated hemoglobin, HDL = high density lipoprotein, hsCRP = high sensitivity C-reactive protein.

## RESULTS

### DunedinPACNI: a Brain MRI Measure of Longitudinal Aging

We trained an elastic net regression model to predict the longitudinal Pace of Aging measure using T1-weighted MRI scans collected in a subsample of 860 Dunedin Study members when they were 45 years old. This subsample maintains the population-representativeness of the full cohort (see **Supplemental Figures S1-S2**). Specifically, the elastic net regression model used 315 MRI-derived structural measures for each Study member including regional cortical thickness, surface area, gray matter volume, and gray-white signal intensity ratio as well as subcortical gray matter and ventricular volumes^37^. We performed 10-fold cross-validation to identify optimal tuning parameters^38^. This optimized model was used to create DunedinPACNI.

The in-sample correlation between DunedinPACNI and the longitudinal Pace of Aging was r=0.60 (**Figure 2A**). We performed a cross-validation analysis by splitting the sample into training and testing subsets 100 different times. Each time, we used 90% of the sample for training and held out the remaining 10% for testing. Across all 100 different splits, the average correlation between DunedinPACNI and Pace of Aging in the testing sample was r=0.42. This prediction accuracy is in line with next-generation epigenetic aging biomarkers^15,20,39^. Of note, for both DunedinPACNI and the longitudinal Pace of Aging, higher scores indicate *faster* aging. DunedinPACNI is a quantitative variable that indexes relative differences between people without clear units. For this reason, we restrict interpretation to directional comparisons within studies. Associations between faster DunedinPACNI scores and measures of physical functioning, cognitive functioning, and facial aging were similar to those previously observed with the Pace of Aging^2^. We controlled for sex in all analyses presented in this manuscript. We further covaried for age in all analyses except those conducted within the Dunedin Study where participants are the same chronological age. Specifically, DunedinPACNI effect sizes for 12 out of the 15 measures were within the 95% confidence intervals of the original Pace of Aging (**Figure 2B**, full results in **Supplemental Table S1**). This was expected given the high internal correlation between DunedinPACNI and Pace of Aging. Dunedin Study members with faster DunedinPACNI scores had worse balance, slower gait, weaker lower- and upper-body strength, and poorer coordination; they also reported worse health and more physical limitations; performed more poorly on tests of cognitive functioning; experienced greater childhood-to-adulthood cognitive decline; and looked older. These results indicate that DunedinPACNI accurately estimates the longitudinal Pace of Aging within the Dunedin Study dataset.

**Figure 2.**
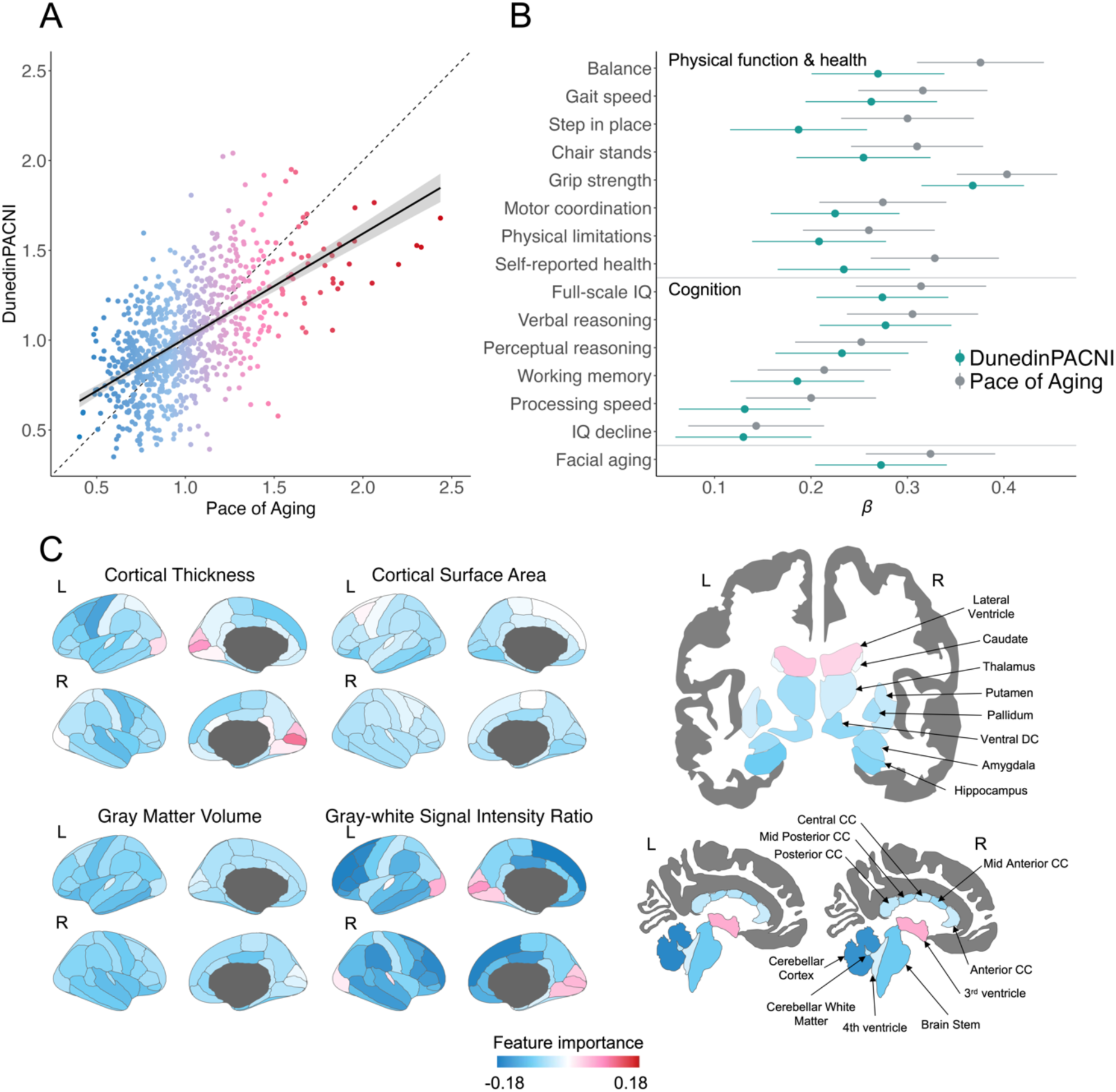
DunedinPACNI model validation and feature importance. **A.** In-sample correlation between Pace of Aging and DunedinPACNI. Warmer colors represent a faster Pace of Aging and cooler colors represent a slower Pace of Aging. **B.** Comparison of absolute effect sizes for associations between DunedinPACNI and Pace of Aging with physical functioning, cognition, and subjective aging measures within the Dunedin Study. Error bars represent the 95% confidence interval. **C.** Covariance between MRI-derived brain features and Pace of Aging. Of the 315 brain features used in model training, 216 were set equal to 0 due to the high correlation between brain measures in order to reduce overfitting. The 99 features included in the final model are visualized in **Supplemental Figure S3.** Warmer colors represent features that positively predicted DunedinPACNI scores (i.e., larger value indicates faster aging) while cooler colors represent features that negatively predicted DunedinPACNI scores (i.e., larger value indicates slower aging). Features that did not contribute to the accuracy of DunedinPACNI predictions are gray. Abbreviations: CC = corpus callosum, DC = diencephalon, L = left, R = right, IQ = Intelligence Quotient.

### DunedinPACNI Reflects Canonical Patterns of Brain Aging

The optimized model used to derive DunedinPACNI included 99 regional brain measures. Due to difficulties in interpreting multivariable model coefficients, we used the Haufe transformation to estimate feature importance scores from the covariance between each brain measure and the Pace of Aging^40^. Given that many of the MRI-derived measures are highly correlated, our elastic net model reduced overfitting by setting the weights for many of them to 0 (visualized in **Supplemental Figure S3**). Faster Pace of Aging covaried with thinner cortex, smaller cortical surface area, smaller cortical gray matter volume, lower cortical gray-white intensity ratio, smaller subcortical gray matter volumes, and larger ventricular volumes (**Figure 2C**). We also observed positive covariance between calcarine cortical thickness and gray-white signal intensity ratio, though not calcarine gray matter volume. This is likely due to known aging-related effects on gray and white matter signal intensity that have been previously demonstrated^41–44^. These structural features overlap with the MRI signatures of both normal brain aging and neurodegenerative diseases^45–49^, suggesting that faster DunedinPACNI reflects, at least in part, canonical patterns of brain aging.

### DunedinPACNI has Excellent Test-Retest Reliability

If DunedinPACNI is to be used as a measure of aging, it must exhibit sufficient measurement reliability when exported to novel datasets. We used test-retest MRI data (N=45) from the Human Connectome Project^50^ to estimate the reliability of DunedinPACNI. The test-retest reliability was excellent (ICC=0.94, 95% CI: [0.89-0.97], **Supplemental Figure S4**).

### DunedinPACNI is Associated with Worse Cognitive Functioning

Having established both internal validity and test-retest reliability, we sought to examine whether DunedinPACNI generalizes to novel datasets to detect aging-related outcomes. Specifically, we first tested for associations with cognitive impairment in ADNI and cognitive functioning in UK Biobank. We generated DunedinPACNI scores from T1-weighted MRI scans collected in 1,737 ADNI participants (mean age=74.3 SD=7.2; range: 52-97 years) and 42,583 UK Biobank participants (mean age=64.4, SD=7.7; range: 44-82 years). In ADNI, participants with faster DunedinPACNI showed greater impairment on mental status exams used to screen for dementia as well as tests of memory, psychomotor speed, and executive functions; they also reported more impairment in cognitively demanding activities of daily living such as maintaining finances or preparing a meal (**Figure 3A**). Absolute standardized effect sizes across all cognitive measures in ADNI ranged from β=0.18 to 0.39 (all p-values<0.001, full results in **Supplemental Table S2**). Similarly, UK Biobank participants with faster DunedinPACNI performed more poorly on tests of executive functions and psychomotor speed (**Figure 3B**). Absolute standardized effect sizes across all cognitive measures in UK Biobank ranged from β=0.05 to 0.17 (all p-values<0.001, full results in **Supplemental Table S3**). Associations in UK Biobank were not driven by individuals with early cognitive decline or *APOE* ε4 homozygotes, who show earlier onset of cognitive decline^51^ (**Supplemental Figure S5**, **Supplemental Table S4**). These, and all subsequent analyses presented in this manuscript, control for sex and chronological age.

**Figure 3.**
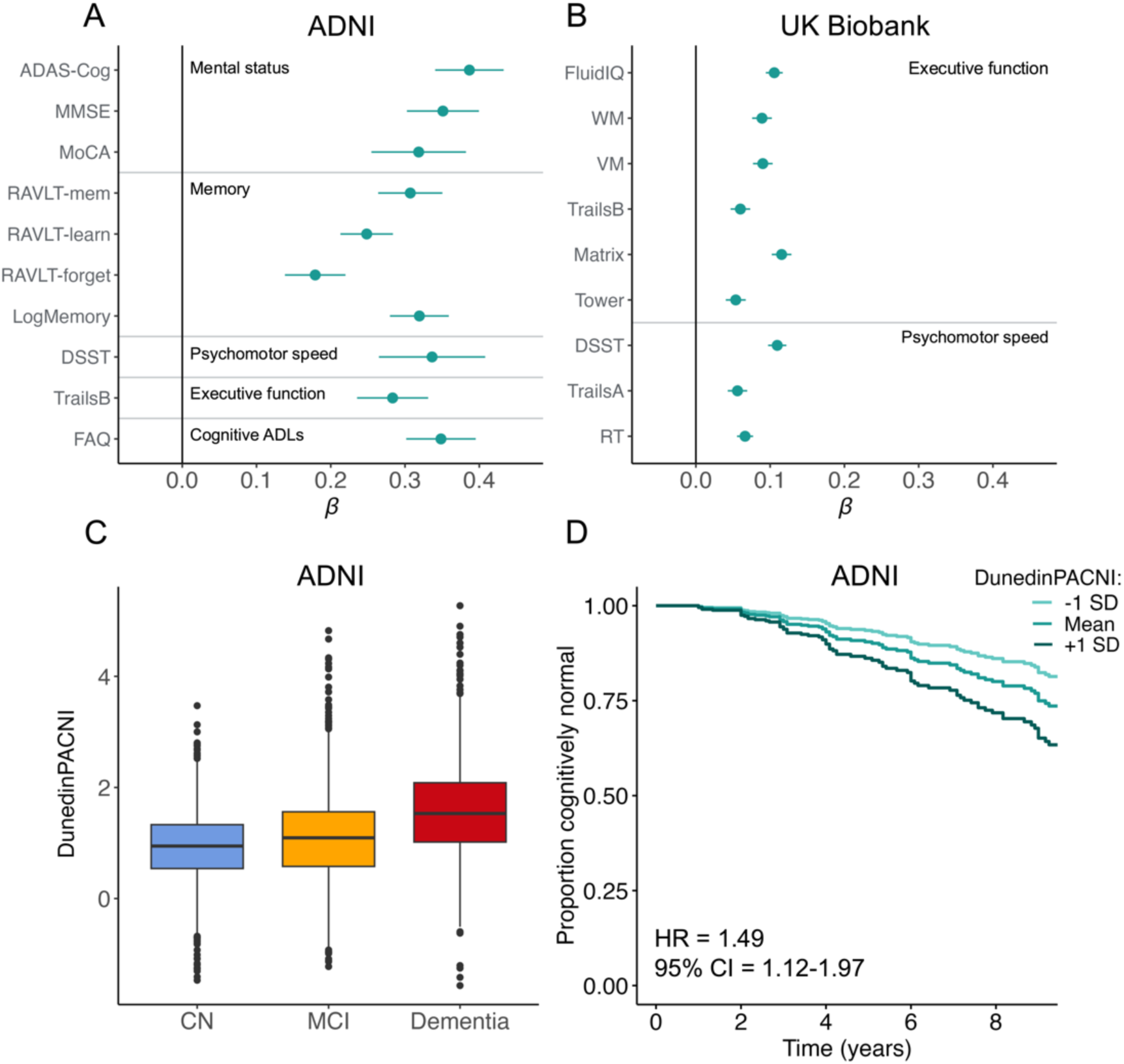
DunedinPACNI predicts cognition, cognitive impairment, and conversion to dementia. Cross-sectional associations between DunedinPACNI and cognitive test scores in A. ADNI and B. the UK Biobank. We visualize absolute effect sizes to aid visual comparison and clarity (see supplemental Tables S2 and S3 for raw effect sizes). Error bars represent the 95% confidence interval. C. Group differences in DunedinPACNI scores amongst ADNI participants according to cognitive status at scanning. Center lines represent the median. Lower and upper hinges represent the 25th and 75th percentiles. Whiskers extend 1.5 times the inter-quartile range from the hinges. Data beyond the whiskers are plotted as individual outliers. D. Survival curve of the relative proportion of cognitively normal ADNI participants at baseline who remained cognitively normal during the follow-up window grouped by slow, average, and fast baseline DunedinPACNI scores. Note that although the maximum follow-up length is 16 years, we have chosen to visualize only 9 years of follow-up due to high amounts of censoring after 9 years. A plot with the full 16 years of follow-up and points marking censoring is presented **in Supplemental Figure S6**. Abbreviations: ADAS-Cog = Alzheimer’s Disease Assessment Scale – Cognitive Subscale 13, ADNI = Alzheimer’s Disease Neuroimaging Initiative, CI = confidence interval, CN = cognitively normal, DSST = Digit Symbol Substitution Task, FAQ = Functional Activities Questionnaire, HR = hazard ratio, IQ = intelligence quotient, LogMemory = Logical Memory, Matrix = Matrix Pattern Completion, MCI = mild cognitive impairment, MMSE = Mini-Mental State Exam, MoCA = Montreal Cognitive Assessment, RT = Reaction Time, RAVLT = Rey Auditory Visual Learning Test, SD = standard deviation, Tower = Tower Rearranging, TrailsA = Trail Making Test Part A, TrailsB = Trail Making Test Part B, VM = Visual Memory, WM = Working Memory.

### DunedinPACNI Predicts Cognitive Decline and Dementia Conversion

We next tested if DunedinPACNI differentiates between normal and clinically impaired cognitive functioning in ADNI (**Figure 3C**). Participants with mild cognitive impairment (MCI) had faster DunedinPACNI compared to cognitively normal participants (β=0.27, p<0.001, 95% CI: [0.18, 0.35]). Participants with dementia had faster DunedinPACNI than both participants with MCI (β=0.54, p<0.001, 95% CI: [0.43, 0.65] and cognitively normal participants (β=0.81 p<0.001, 95% CI: [0.69, 0.92]).

We further tested whether DunedinPACNI predicts future cognitive decline among people without cognitive impairment. Specifically, we analyzed a subsample of 624 ADNI participants who were cognitively normal at the time of their first scan (mean age = 72.4 years, SD = 6.3 years; range = 52.7-89.9 years), 112 of whom converted to either MCI or dementia during up to 16-years of follow-up (mean follow-up = 4.90 years). Cognitively normal participants with faster DunedinPACNI at baseline were more likely to develop MCI or dementia and to do so earlier during the follow-up window (HR=1.49, p=0.005, 95% CI: [1.12, 1.97]; **Figure 3D**), meaning those in the top 10% had a 61% increased risk of developing MCI or dementia compared to participants with average DunedinPACNI. We conducted a similar analysis among the 701 participants who were diagnosed with MCI at the time of their first scan (mean age = 72.8 years, SD = 7.3 years, range = 55.0-88.8 years), 271 of whom converted to dementia during up to 16 years of follow-up (mean follow-up = 4.10 years). MCI participants with faster DunedinPACNI at baseline were more likely to convert to dementia (HR=1.44, p<0.001, 95% CI: [1.26, 1.65]). These effect sizes were similar when controlling for number of *APOE* ε4 alleles, a well-established genetic risk allele for sporadic, late-onset Alzheimer’s disease (baseline cognitively normal: HR=1.49, p=0.005, 95% CI: [1.13, 1.96]; baseline MCI: HR=1.42, p<0.001, 95% CI: [1.23, 1.62]). Because only a very small number of UK Biobank participants with MRI data received diagnoses of dementia during follow-up observation (N=73), we were underpowered to report parallel results in this dataset.

### DunedinPACNI Predicts Accelerated Brain Atrophy

As an estimate of how fast a person is aging, DunedinPACNI should reflect longitudinal trajectories of brain decline^33^. We tested whether faster baseline DunedinPACNI predicted accelerated hippocampal atrophy, which is an established risk factor for cognitive decline and dementia onset in older adults^52^. Specifically, we computed longitudinal trajectories of hippocampal atrophy among 1,302 ADNI participants who had MRI data at multiple time points (average number of scans=4.4, range=2 to 13 scans) as well as 4,628 UK Biobank participants who had MRI data at two timepoints. Participants with faster baseline DunedinPACNI exhibited accelerated hippocampal atrophy, in both ADNI (β=-0.15, p<0.001, 95% CI: [-0.21, -0.10]; **Figure 4B**) and the UK Biobank (β=-0.09, p<0.001, 95% CI: [-0.12, -0.05]; **Figure 4B**). This result was consistent while controlling for number of *APOE* ε4 alleles (**Supplemental Table S4**).

**Figure 4.**
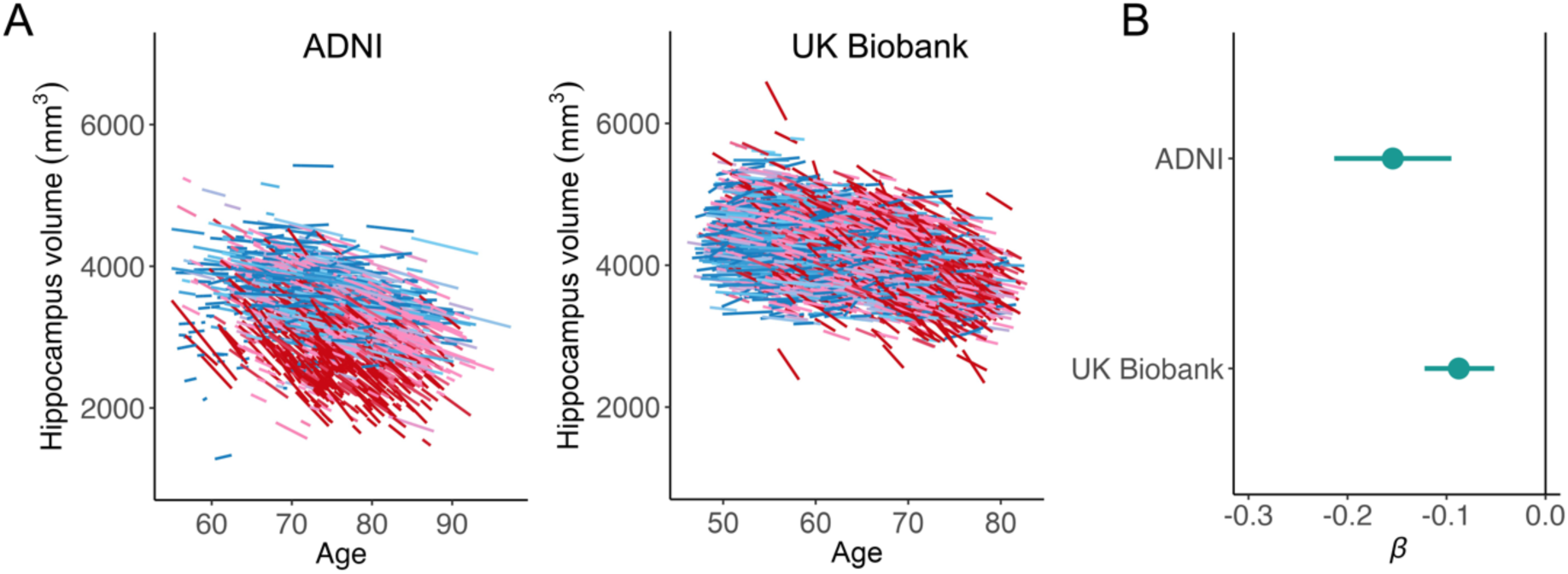
DunedinPACNI predicts accelerated hippocampal atrophy. **A.** Individualized trajectories of hippocampal atrophy in ADNI and UK Biobank. Warmer colors represent accelerated atrophy. **B.** Forest plot of associations between baseline DunedinPACNI scores and accelerated hippocampal atrophy in ADNI and UK Biobank. Error bars represent 95% confidence intervals and effect sizes. Abbreviations: ADNI = Alzheimer’s Disease Neuroimaging Initiative, mm^3^ = cubic millimeters.

### DunedinPACNI Predicts Frailty, Poor Health, Chronic Disease, and Mortality

As a measure of aging derived from longitudinal assessments of multiple biomarkers, DunedinPACNI should capture instances of declining health across all organ systems, not just the brain. To test this hypothesis, we used the UK Biobank to map DunedinPACNI scores onto measures of frailty, subjective overall health, incident aging-related chronic diseases, and all-cause mortality.

We used the Fried Frailty Index to quantify the degree of vulnerability to common stressors associated with aging-related decline in energy reserves and functioning. When treating index scores as a continuous measure ranging from 0 to 5 with higher scores indicating greater frailty^53,54^, we found that participants with faster DunedinPACNI were more frail (N=42,583; β=0.17, p<0.001, 95% CI:[0.16, 0.18]). Participants with faster DunedinPACNI also self-reported poorer overall health (N=42,235; β=-0.17, p<0.001, 95% CI:[- 0.18, -0.16]; **Figure 5A**), which predicts mortality even independently of objective health measures^55^. These associations were not driven by individuals with early cognitive decline or high genetic risk for Alzheimer’s disease (**Supplemental Figure S5**, **Supplemental Table S4**).

**Figure 5.**
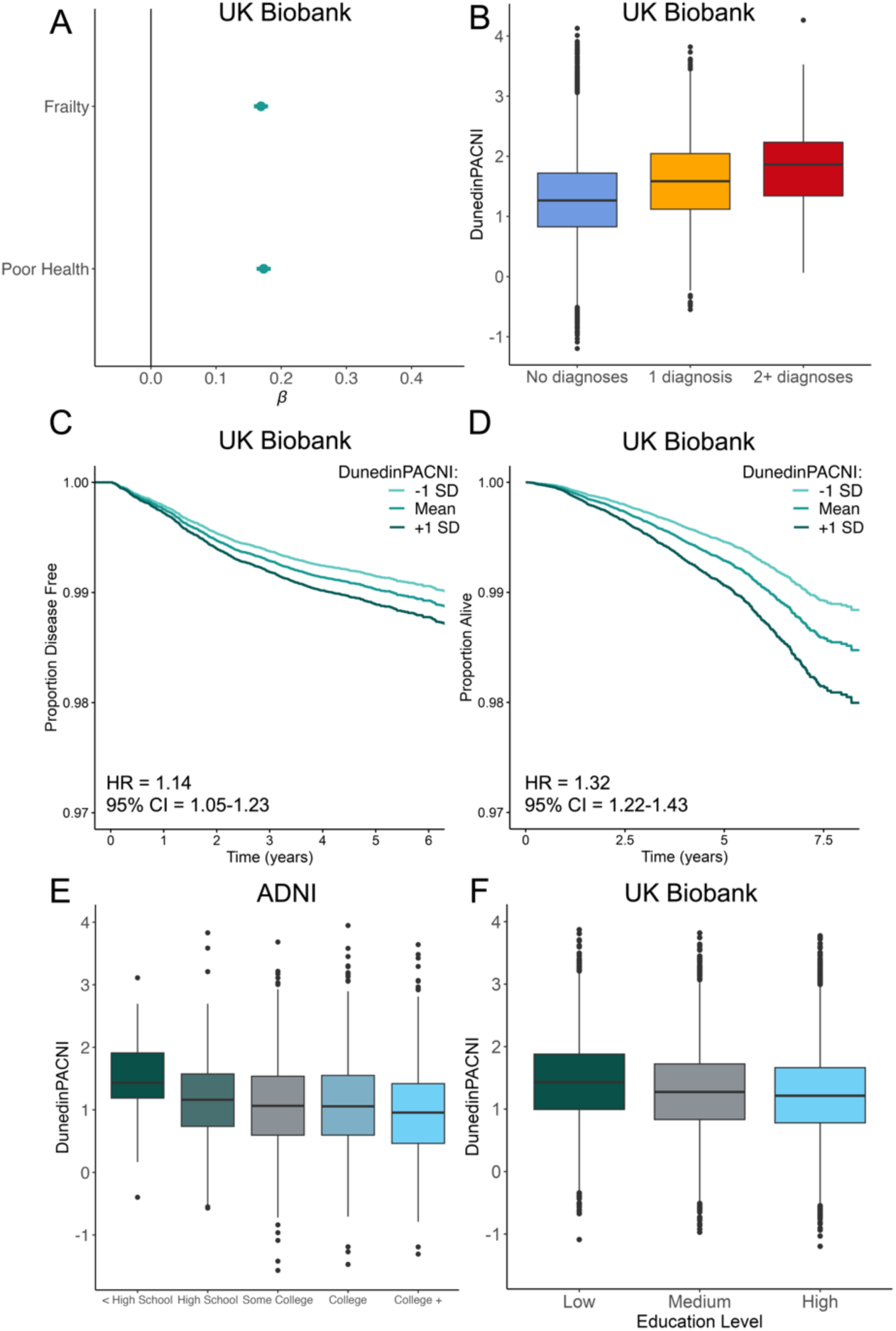
DunedinPACNI predicts frailty, poor health, multimorbidity, future chronic diseases, and mortality, and reflects social gradients of health inequities. **A.** Forest plot of absolute associations between DunedinPACNI and self-rated health and frailty in the UK Biobank. Error bars represent 95% confidence intervals. **B**. Group differences in DunedinPACNI scores according to lifetime number of aging-related chronic disease diagnoses including myocardial infarction, chronic obstructive pulmonary disease, dementia, and stroke in the UK Biobank. **C**. Survival curve of the relative proportion of disease-free UK Biobank participants at time of MRI who remained disease-free during the follow-up window, grouped by slow, average, and fast baseline DunedinPACNI scores. Of note, we excluded participants who had chronic disease prior to scanning from this analysis. **D.** Survival curve of the relative proportion of UK Biobank participants who remained alive during the follow-up window grouped by baseline DunedinPACNI scores. **E.** Group differences in DunedinPACNI according to education level among ADNI participants. **F.** Group differences in DunedinPACNI according to education level in the UK Biobank. For boxplots in **B**, **E**, and **F**, Center lines represent the median. Lower and upper hinges represent the 25^th^ and 75^th^ percentiles. Whiskers extend 1.5 times the inter-quartile range from the hinges. Data beyond the whiskers are plotted as individual outliers. Abbreviations: ADNI = Alzheimer’s Disease Neuroimaging Initiative, CI = confidence interval, HR = hazard ratio, SD = standard deviation.

Similar patterns emerged when considering clinical diagnoses of chronic aging-related diseases including myocardial infarction, chronic obstructive pulmonary disease, dementia, and stroke. Participants with a lifetime prevalence of one of these chronic diseases had faster DunedinPACNI compared to those reporting no diagnoses (β=0.19, p<0.001, 95% CI: [0.16, 0.23]). Participants with a lifetime prevalence of two or more chronic diseases had faster DunedinPACNI than those with a single chronic disease (β=0.25, p<0.001, 95% CI: [0.12, 0.38] and those with no chronic disease (β=0.44, p<0.001, 95% CI: [0.31, 0.57]; **Figure 5B**).

Extending beyond contemporaneous associations, we assessed whether faster DunedinPACNI at baseline predicted future myocardial infarction, chronic obstructive pulmonary disease, dementia, or stroke in UK Biobank participants who were diagnosis-free at the time of scanning (N=40,753). 827 participants reported a new diagnosis of at least one of these aging-related chronic diseases over a maximum follow-up period of 9.7 years after scanning (i.e., baseline). Consistent with the above contemporaneous associations, healthy participants with faster DunedinPACNI at baseline were more likely to be later diagnosed with chronic aging-related diseases (HR=1.14, p<0.001, 95% CI: [1.05, 1.23]; **Figure 5C**), meaning those in the top 10% had an 18% or greater increased risk for developing a chronic disease compared to participants with average DunedinPACNI. These associations were not driven by individuals with early cognitive decline or high genetic risk for Alzheimer’s disease (**Supplemental Figure S5**, **Supplemental Table S4**).

Given the increased mortality rates amongst people with chronic aging-related diseases, we asked if baseline DunedinPACNI scores predicted all-cause mortality. Of the 42,583 UK Biobank participants included in our dataset, 757 died over the follow-up period after their baseline MRI scan. Participants with faster baseline DunedinPACNI scores died earlier (HR=1.32, p<0.001, 95% CI: [1.22, 1.43]; **Figure 5D**), meaning those in the top 10% were at least 41% more likely to die compared to participants with average DunedinPACNI. These associations were not driven by individuals with early cognitive decline or high genetic risk for Alzheimer’s disease (**Supplemental Figure S5**, **Supplemental Table S4**). Taken together, these findings suggest that DunedinPACNI is useful for gauging general physical health and assessing risk for future chronic disease and death.

### DunedinPACNI Reflects Social Gradients of Health Inequities

People who are less advantaged in their socioeconomic position experience a wide range of chronic diseases and earlier mortality^56–58^, and DunedinPACNI should reflect such gradients of health inequities. We used information about educational attainment and income to test this prediction. Faster DunedinPACNI was observed for participants who either had fewer years of formal education (ADNI: β=- 0.10, p<0.001, 95% CI: [-0.15, -0.05]; UK Biobank: β=-0.06, p<0.001, 95% CI: [-0.07, -0.05]) or lower income (UK Biobank: β=-0.09, p<0.001, 95% CI: [-0.10, -0.08]), reflecting the expected socioeconomic health gradient (**Figure 5E-F**). These associations were not driven by individuals with early cognitive decline or high genetic risk for Alzheimer’s disease (**Supplemental Figure S5**, **Supplemental Table S4**).

### DunedinPACNI generalizes to a Latin American sample

Brain-based predictive algorithms often fail when applied in groups that demographically differ from the training sample^59^. The large majority of cognitive and brain aging studies are collected in high-income countries in North America and Europe, which may limit the generalizability of brain-based models to people in low- and middle-income countries^60–62^. Latin Americans are underrepresented in biomedical research^61,63^ and may experience distinct social and environmental influences on brain aging compared to people in high-income countries in North America and Europe^61^. To test the generalizability of DunedinPACNI among Latin Americans, we calculated DunedinPACNI scores in a sample of 369 adults from Argentina, Chile, Colombia, Mexico, and Peru from the BrainLat dataset^64^. 162 BrainLat participants were diagnosed with Alzheimer’s disease (AD), 84 were diagnosed with behavior-variant frontotemporal dementia (FTD), and 123 were healthy controls. When controlling for age and sex, BrainLat participants with AD and FTD had faster DunedinPACNI scores compared to healthy controls (AD: β=0.70, p<0.001, 95% CI: [0.48, 0.91]; FTD: β=0.79, p<0.001, 95% CI: [0.55, 1.03]; **Figure 6A**). Of note, these effect sizes are comparable to the difference between dementia and cognitively normal participants in ADNI (**Figure 6B**). In addition, 191 BrainLat participants also completed the Montreal Cognitive Assessment (MoCA). Among this subset, BrainLat participants with faster DunedinPACNI scores had poorer scores on the MoCA (β=- 0.35, p<0.001, 95% CI: [-0.49, -0.20]). Of note, this effect was similar to the association between DunedinPACNI and MoCA scores in ADNI participants (β=-0.32, p<0.001, 95% CI: [-0.38, -0.25]; **Figure 6C**).

**Figure 6.**
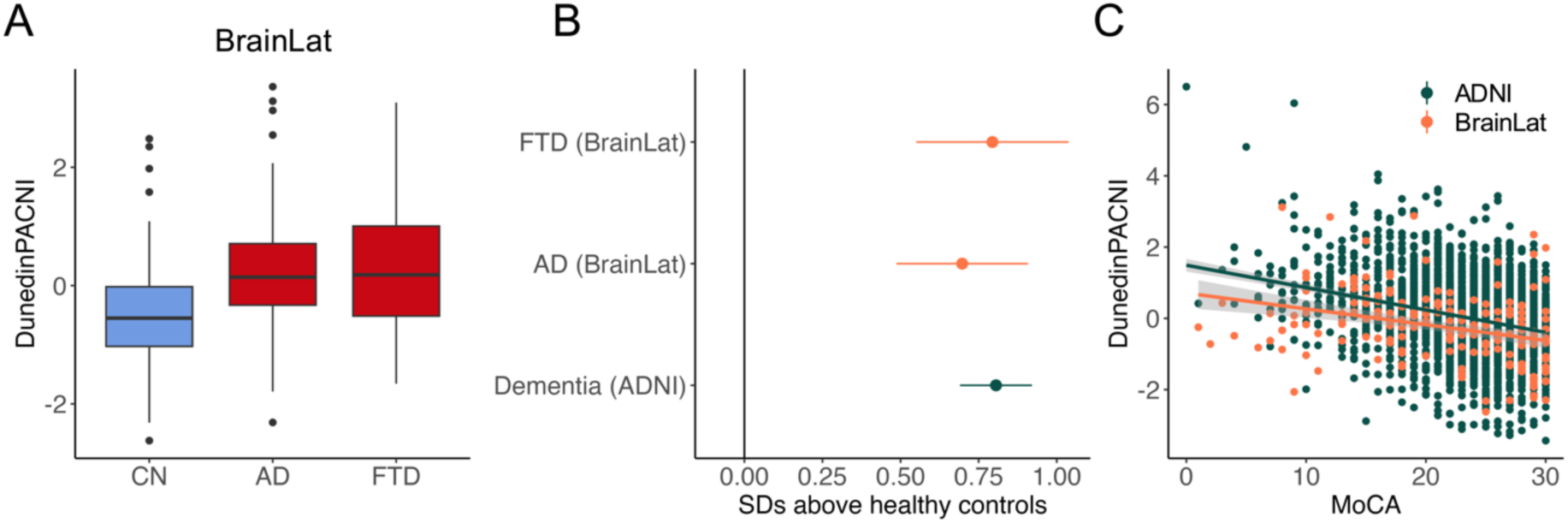
DunedinPACNI is similarly associated with dementia and cognitive impairment in a sample of Latin Americans. **A.** Group differences in DunedinPACNI scores according to cognitive diagnosis in BrainLat. Center lines represent the median. Lower and upper hinges represent the 25^th^ and 75^th^ percentiles. Whiskers extend 1.5 times the inter-quartile range from the hinges. Data beyond the whiskers are plotted as individual outliers. DunedinPACNI scores are residualized for age and sex. **B.** Forest plot of standardized mean differences in DunedinPACNI between dementia and cognitively normal healthy controls in BrainLat (orange) and ADNI (dark green) while controlling for age and sex. DunedinPACNI shows similar accelerated in dementia in a sample of Latin Americans (BrainLat) and North Americans (ADNI). Error bars represent 95% confidence intervals. **C.** Scatter plot of associations between MoCA scores and DunedinPACNI in BrainLat (orange) and ADNI (dark green). DunedinPACNI scores are residualized for age and sex. Linear associations between MoCA scores and DunedinPACNI scores are similar in a sample of Latin Americans (BrainLat) and North Americans (ADNI). Abbreviations: AD = Alzheimer’s disease, ADNI = Alzheimer’s Disease Neuroimaging Initiative, CN = cognitively normal, FTD = frontotemporal dementia, MoCA = Montreal Cognitive Assessment, SD = standard deviation.

### DunedinPACNI is Distinct from Brain Age Gap and Commonly Reported Measures of Brain Aging

Lastly, we compared DunedinPACNI with existing approaches for measuring aging using brain MRI data. Specifically, we compared effect sizes for DunedinPACNI from all of the aforementioned analyses in ADNI, UK Biobank, and BrainLat with brain age gap generated using brainageR^35^. We selected this algorithm due to its high accuracy and test-retest reliability compared to other brain age gap algorithms^65^. DunedinPACNI and brain age gap were only modestly correlated (ADNI: r=0.17, p <0.001; UK Biobank: r=0.31, p<0.001; BrainLat: r=0.32, p<0.001; **Supplemental Figure S7**). Compared to brain age gap, the effect sizes for DunedinPACNI were similar or larger across measures of cognitive function, cognitive decline, brain atrophy, frailty, disease risk, mortality, and socioeconomic health gradients (**Figure 7, Supplemental Figure S8-S9,** full results in **Supplemental Tables S2-S3, S6-S11**). Commensurate with the low correlation between these measures, when we included both DunedinPACNI and brain age gap in a single model, each measure explained unique variance in clinical outcomes with only minor reductions in effect sizes. Moreover, using both DunedinPACNI and brain age gap together in a single model generally increased prediction of these outcomes (**Supplemental Figure S8-S9**). For example, the combined hazard ratio of DunedinPACNI and brain age gap predicting mortality risk was 1.50 (95% CI: [1.36, 1.65]), compared to the independent hazard ratios of 1.32 for DunedinPACNI and 1.24 for brain age gap.

**Figure 7.**
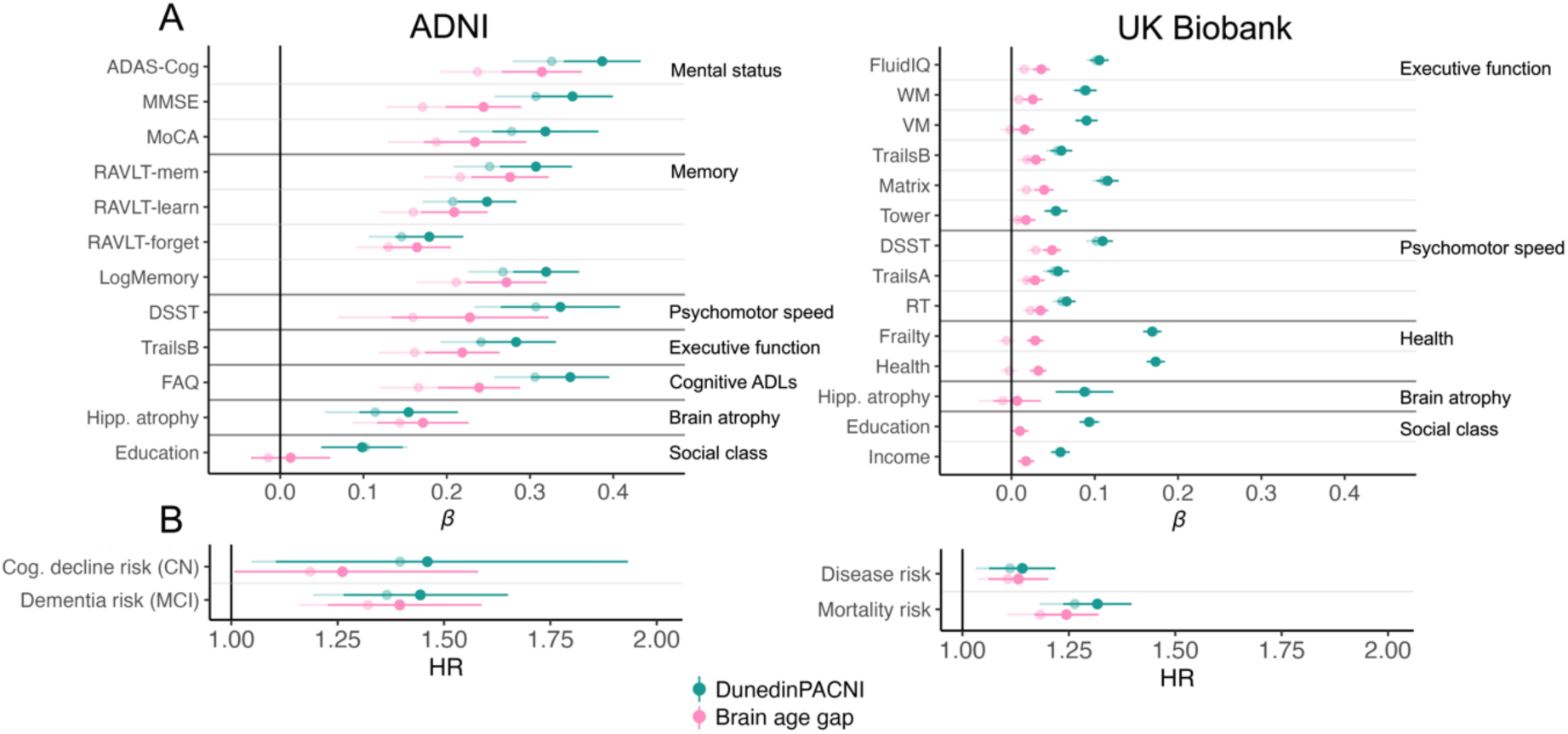
Comparison of DunedinPACNI and brain age gap associations with aging-related phenotypes. **A.** Forest plots of DunedinPACNI and brain age gap absolute effect sizes in ADNI and UK Biobank. **B.** Forest plots of DunedinPACNI and brain age gap hazard ratios in ADNI and UK Biobank. Error bars represent 95% confidence intervals. Lighter shades represent the effect size for each measure while controlling for the other measure (i.e., effect of DunedinPACNI when controlling for brain age gap, and vice versa). Abbreviations: ADAS-Cog = Alzheimer’s Disease Assessment Scale – Cognitive Subscale 13, ADLs = activities of daily living, ADNI = Alzheimer’s Disease Neuroimaging Initiative, CI = confidence interval, CN = cognitively normal, Cog. = cognitive, DSST = Digit Symbol Substitution Task, FAQ = Functional Activities Questionnaire, Hipp. = hippocampal, HR = hazard ratio, IQ = intelligence quotient, LogMemory = Logical Memory, Matrix = Matrix Pattern Completion, MCI = mild cognitive impairment, MMSE = Mini-Mental State Exam, MoCA = Montreal Cognitive Assessment, RT = Reaction Time, RAVLT = Rey Auditory Visual Learning Test, SD = standard deviation, Tower = Tower Rearranging, TrailsA = Trail Making Test Part A, TrailsB = Trail Making Test Part B, VM = Visual Memory, WM = Working Memory.

In addition, we investigated how DunedinPACNI differs from commonly reported MRI-based measures of brain aging, namely hippocampal volume and ventricular volume. To test this, we compared effect sizes for DunedinPACNI with effect sizes for bilateral hippocampal volume and bilateral ventricular volume in UK Biobank and among cognitively normal participants in ADNI. In both datasets, we observed that faster DunedinPACNI was generally more strongly and more consistently associated with poor cognition, poor health, and frailty, as well as risk of dementia, disease, and mortality. Furthermore, DunedinPACNI explained incremental variance in these outcomes over and above hippocampal volume and ventricular volume alone (**Supplemental Figures S10-S12, Supplemental Tables S12-S14**).

## DISCUSSION

DunedinPACNI is an accurate and reliable measure of how fast a person is aging derived from a single brain MRI scan. Using 50,106 scans from people ranging from 22-98 years old across the HCP, ADNI, UK Biobank, and BrainLat cohorts we demonstrate that people with faster DunedinPACNI had not only worse cognitive and brain health (i.e., poorer cognition, faster hippocampal atrophy, and greater dementia risk) but also worse general health (i.e., greater frailty, poorer self-reported health, greater risk of chronic disease and mortality). This indicates that patterns of aging detected during midlife are clinically useful among people in advanced age, including people with neurodegenerative disease. Furthermore, DunedinPACNI showed evidence of generalization to a sample of Latin American adults with and without dementia. Across all analyses, the effect sizes for DunedinPACNI were similar or larger than the effect sizes for brain age gap, an existing age-deviation measure derived from the same structural MRI data. Moreover, DunedinPACNI and brain age gap were only weakly correlated, and DunedinPACNI accounted for incremental variance in aging-related health outcomes. While weak correlations between neuroimaging-based measures of aging may appear surprising, they mirror findings that different epigenetic clocks are also weakly correlated, and multiple clocks are useful for predicting disease and death^66,67^. Aging biomarkers trained in midlife samples may more closely detect the earlier stages of aging compared to aging biomarkers trained in samples of advanced age. This may partially explain the modest correlations observed between aging biomarkers. Aging remains a construct in search of measurement tools^4,5^, and DunedinPACNI represents a “next-generation” measure of aging that is distinct from existing approaches.

DunedinPACNI is not without limitations. First, the Dunedin Study, ADNI, and UK Biobank consist of data collected primarily from participants of European ancestry. Furthermore, ADNI and UK Biobank over-sample participants from higher socioeconomic backgrounds^68^. There is growing awareness that the lack of representativeness in neuroimaging research may hinder clinical translation^69,70^, including of brain-based predictive models^59^. To address this, we replicated associations with DunedinPACNI among a sample of Latin Americans, as well as UK Biobank participants who are low-income or non-White. (**Supplemental Tables S15-S16**). Additionally, our findings in ADNI, UK Biobank, and BrainLat demonstrate that DunedinPACNI generalizes well to older adults. This suggests that early differences in aging can be detected in brain MRI measures collected at age 45 that are clinically useful in people who are decades older. At the same time, it is also possible that an analogous aging biomarker trained in older adults may generalize more readily to other samples of older adults. A priority for future work is to further evaluate the generalizability of DunedinPACNI to people of diverse demographics and backgrounds. Second, DunedinPACNI only uses structural brain measures derived from a T1-weighted MRI scan. We chose this strategy because these scans are collected in nearly every MRI study, thereby maximizing the potential adoption of DunedinPACNI. It is possible that the performance accuracy reported here could be improved by including additional structural and/or functional MRI measures (e.g., white matter microstructural integrity from diffusion weighted images, BOLD signal from T2*-weighted images). Relatedly, while elastic net is widely used in biological age measurement^11,12,15,16,20^, future research should evaluate whether predictive performance accuracy could be increased by more complex, non-linear models. Third, by design, DunedinPACNI is a measure of the longitudinal rate of the aging of the body derived from a single MRI scan and is not designed to replace longitudinal measurement of brain aging through repeated MRI assessments. Fourth, DunedinPACNI is estimated through observed correlations between measures of brain structure and longitudinal aging, which could reflect multiple causal pathways. For example, faster aging of non-brain organs might cause poorer brain health or vice versa. Alternatively, both may be driven by a third factor. Fifth, although we found robust associations with aging phenotypes across both ADNI and UK Biobank, we generally observed larger effect sizes in ADNI. This could suggest that DunedinPACNI is sensitive to dysfunction among patients with neurodegenerative diseases. Further evaluation is needed to establish the degree to which DunedinPACNI is sensitive to individual organ systems. Sixth, DunedinPACNI is currently a quantitative tool for comparison of individuals within datasets but not between datasets. Future research, potentially through data harmonization and normative modeling approaches, should establish normative reference values and ranges that can aid in the clinical interpretation of DunedinPACNI scores. Seventh, as is true for all MRI-based measurements, the derivation of DunedinPACNI may be compromised in low quality scans such as those with high levels of head motion. In the analyses presented here, we followed standard practice by excluding low quality scans (**Supplemental Figures S13-S16**). Future research should adopt methods to improve data quality in high-motion participants to help maximize generalizability^71,72^.

Currently, DunedinPACNI and general aging biomarkers are research tools that require further validation before potential translation to the clinic^5^. Nonetheless, we believe DunedinPACNI and other general aging biomarkers have several promising uses that differ from and are complementary to biomarkers of specific diseases (e.g. hippocampal atrophy and Alzheimer’s disease^73^ or high blood pressure and stroke^74^). For example, general aging biomarkers could help identify a broad risk for many age-related diseases earlier in the lifespan when individuals may benefit most from health and anti-aging intervention^23^. Furthermore, general aging biomarkers could help establish a person’s prognosis after the onset of a specific disease as well as how interventions or insults may change how fast a person is aging^25,75,76^. Lastly, general aging biomarkers are necessary to mechanistically link exposures (e.g., low socioeconomic status) to aging or to link the rate of aging to outcomes such as increased disease risk^4^. While detecting or diagnosing a specific disease requires a proximal, specific biomarker of that disease, general aging biomarkers could be used to measure distal risk for a broad array of diseases. This difference allows for novel clinical applications for aging biomarkers such as DunedinPACNI alongside biomarkers of specific diseases.

Several unique features of the Dunedin Study contribute advantages to DunedinPACNI compared to other aging biomarkers. First, DunedinPACNI was developed in a cohort of people all born in the same year and studied at the same ages throughout their lives, thereby avoiding biases that are introduced by differences in historical exposures across generations and across time. Second, DunedinPACNI is trained on 19 biomarkers that were each assessed over two decades and thus are not influenced by short-term illnesses that can cause aberrant biomarker signals at a single assessment. Third, DunedinPACNI was derived from participants followed from birth to age 45 before the onset of chronic, aging-related diseases that cause divergence from typical trajectories of aging. Other aging biomarkers are typically derived from multiage samples in which many of the older members already have chronic diseases that have altered their body and brain, so those measures may inadvertently repackage disease instead of aging. Fourth, because the Dunedin Study cohort is population-representative with very low attrition and mortality rates, DunedinPACNI does not suffer from oversampling of healthy volunteers, attrition bias (i.e., people with worse health being more likely to drop out), or survivor bias (i.e., people with worse health dying earlier). Indeed, our results align with prior research showing that DunedinPACE, which like DunedinPACNI was trained on the longitudinal Pace of Aging, is associated with dementia, morbidity, and mortality^20,22,23^. Our results, alongside the fast-growing literature on DunedinPACE, suggest that these unique design characteristics of the Dunedin Study make it a powerful training sample for longitudinal aging biomarkers.

The scope of geroscience has rapidly expanded with the proliferation of -omic clocks that can measure how fast people age^10^. DunedinPACNI is poised to further this growth by allowing individual differences in the rate of longitudinal aging to be estimated from a single noninvasive MRI scan that can be collected in just a few minutes. Indeed, the requisite MRI data to estimate DunedinPACNI have already been collected in many psychiatric, neurologic, and brain-health cohorts, from tens of thousands of research participants across the lifespan and around the world. DunedinPACNI offers an opportunity to enrich such studies and deepen understanding of the causes of individual differences in the rate of longitudinal aging, including genetics^77^, childhood adversities^78^, environmental exposures (e.g., lead^79,80^), and lifestyle factors (e.g., physical inactivity, social isolation^81^). DunedinPACNI may also be adopted as a surrogate endpoint to accelerate our ability to develop, prioritize, and evaluate potential anti-aging interventions that slow aging and prevent disease^82,83^. The algorithm for DunedinPACNI will be made publicly available to the research community to facilitate these and other future research directions.

## METHODS

The premise and analysis plan for this study were pre-registered (link: https://rb.gy/b9x4u6). All analyses and code were checked for accuracy by an independent analyst. Analyses were conducted on data collected through the Dunedin Study, Human Connectome Project, Alzheimer’s Disease Neuroimaging Initiative, UK Biobank, and BrainLat. Details for each study and dataset are described below.

## DATA SOURCES

### Dunedin Study

Participants are members of the Dunedin Study, a longitudinal investigation of health and behavior in a representative birth cohort. The 1,037 participants (91% of eligible births, 48% female) were all people born between April 1972 and March 1973 in Dunedin, New Zealand, who were residents in the province and who participated in the first assessment at age 3 years^19^. The cohort represented the full range of socioeconomic status in the general population of New Zealand’s South Island and, as adults, matches the New Zealand National Health and Nutrition Survey on key adult health indicators (e.g., body mass index, smoking, and general practitioner visits) and the New Zealand Census of citizens of the same age on educational attainment^19,84^. Study participants are primarily of New Zealand European ethnicity; 8.6% reported Māori ethnicity at age 45.

General assessments were performed at birth as well as ages 3, 5, 7, 9, 11, 13, 15, 18, 21, 26, 32, and 38 years; and, most recently (completed April 2019), at age 45 years, when 938 of the 997 living Study members (94.1%) participated. At each assessment, Study members were brought to the Dunedin Study Research Unit at the University of Otago for interviews and examinations. In addition, staff provided standardized ratings, informant questionnaires were sent to people who the Study members nominated as people who knew them well, and administrative records were searched. The Dunedin Study was approved by the University of Otago Ethics Committee and Study members gave written informed consent before participating.

#### MRI

As a component of the age 45 assessments, Study members were scanned using a Siemens MAGNETOM Skyra (Siemens Healthcare GmbH) 3T scanner equipped with a 64-channel head/neck coil at the Pacific Radiology Group imaging center in Dunedin, New Zealand. High resolution T1-weighted images were obtained using an MP-RAGE sequence with the following parameters: TR=2400 ms; TE=1.98 ms; 208 sagittal slices; flip angle, 9°; FOV, 224 mm; matrix =256×256; slice thickness=0.9 mm with no gap (voxel size 0.9×0.875×0.875 mm); and total scan time=6 min and 52 s. 3D fluid-attenuated inversion recovery (FLAIR) images were obtained with the following parameters: TR=8000 ms; TE=399 ms; 160 sagittal slices; FOV=240 mm; matrix=232×256; slice thickness=1.2 mm (voxel size 0.9×0.9×1.2 mm); and total scan time=5 min and 38 s. Additionally, a gradient-echo field map was acquired with the following parameters: TR=712 ms; TE=4.92 and 7.38 ms; 72 axial slices; FOV=200 mm; matrix=100×100; slice thickness=2.0 mm (voxel size 2 mm isotropic); and total scan time=2 min and 25 s. Of the 938 Study members seen at Phase 45, 63 declined to participate in MRI scanning, meaning 875 Study members completed the MRI scanning protocol. Scanned Study members did not differ significantly from other living participants in terms of childhood neurocognitive functioning or childhood SES (see attrition analysis in the **Supplemental Figures S1-S2**). Of these 875 Study members for whom data was available, 4 were excluded due to major incidental findings or previous injuries (e.g. large tumors or extensive damage to the brain/skull), 9 due to missing FLAIR or field map scans, 1 due to poor surface mapping yielding, and 1 due to missing the Pace of Aging variable. This yielded a final training sample of 860 Study members (see **Supplemental Figure S13** for inclusion details).

Structural MRI data were processed using FreeSurfer version 6.0^37^. Specifically, T1-weighted images were processed and refined with 3D FLAIR images using the recon-all pipeline.

#### Pace of Aging

Participants’ pace of biological aging was measured as changes in 19 biomarkers of cohort members’ cardiovascular, metabolic, pulmonary, kidney, immune and dental systems across ages 26, 32, 38, and 45 years. This measure quantifies participants’ rate of aging in year-equivalent units of physiological decline per chronological year. The average participant experienced 1 year of physiological decline per year, a mean (SD) Pace of Aging of 1 (0.3)^2^. See the Statistical Analysis section for more details.

#### Physical Functioning

One-legged balance was measured using the Unipedal Stance Test as the maximum time achieved across three trials of the test with eyes closed^87–89^. Gait speed (meters per second) was assessed with the 6-m-long GAITRite Electronic Walkway (CIR Systems, Inc) with 2-m acceleration and 2-m deceleration before and after the walkway, respectively. Gait speed was assessed under 3 walking conditions: usual gait speed (walk at a normal pace from a standing start, measured as a mean of 2 walks) and 2 challenge paradigms, dual-task gait speed (walk at a normal pace while reciting alternate letters of the alphabet out loud, starting with the letter “A,” measured as a mean of 2 walks) and maximum gait speed (walk as fast as safely possible, measured as a mean of 3 walks). Gait speed was correlated across the 3 walk conditions^90^. To increase reliability and take advantage of the variation in all 3 walk conditions (usual gait and the 2 challenge paradigms), we calculated the mean of the 3 highly correlated individual walk conditions to generate our primary measure of composite gait speed. The step in place test was measured as the number of times the right knee was lifted to mid-thigh height (measured as the height half-way between the knee cap and the iliac crest) in 2 minutes at a self-directed pace^91^. Chair rises were measured as the number of stands with no hands completed in 30 seconds from a seated position^92^. Handgrip strength was measured for each hand (elbow held at 90°, upper arm held tight against the trunk) as the maximum value achieved across three trials using a Jamar digital dynamometer^91,93^. Analyses using handgrip strength controlled for BMI. Visual-motor coordination was measured as the time to completion of the Grooved Pegboard Test. Scores were reversed so that higher values corresponded to better performance. Physical limitations were measured with the RAND 36-Item Health Survey 1.0 physical functioning scale. Responses (“limited a lot”, “limited a little”, “not limited at all”) assessed difficulty with completing various activities (e.g., climbing several flights of stairs, walking more than 1 km, participating in strenuous sports). Scores were reversed to reflect physical limitations so that a high score indicates more limitations.

#### Subjective Health and age appearance

We obtained reports about Study members’ health and age appearance from three sources: self-reports, informant impressions, and staff impressions. We obtained reports about Study members’ age appearance from three sources: self-reports, informant impressions, and staff impressions. *Self-reports* – We asked the Study members about their own impressions of how old they looked, “Do you think you LOOK older, younger, or about your actual age?” Response options were younger than their age, about their actual age, or older than their age. We also asked Study members to rate their age perceptions in years, “How old do you feel?” *Informant impressions* - Informants who knew a Study member well (94% response rate) were asked: “Compared to others their age, do you think he/she (the Study member) looks younger or older than others their age? Response options were: “much younger”, “a bit younger”, “about the same”, “a bit older”, or “much older”. *Staff impressions* - Four members of the Dunedin Study Unit staff completed a brief questionnaire describing each study member. To assess age appearance, staff used a 7-item scale to assign a “relative age” to each Study member (1=young looking, 7=old looking). Correlations between self-, informant-, and staff-ratings ranged from 0.34–0.52. All reporters rated the Study member’s general health using the following response options: excellent, very good, good, fair, or poor. Correlations between self-, informant-, and staff-ratings ranged from *r*=*0*.48–0.55.

#### Cognitive Functioning

The Wechsler Adult Intelligence Scale-IV (WAIS-IV) was administered at age 45, yielding adult IQ. In addition to full-scale IQ, the WAIS-IV yields indexes of four specific cognitive function domains: Processing Speed, Working Memory, Perceptual Reasoning, and Verbal Comprehension. The Wechsler Intelligence Scale for Children–Revised (WISC–R) was administered at ages 7, 9, and 11 years. To increase baseline reliability the three scores were averaged yielding childhood IQ. We measured cognitive decline by studying adult IQ scores after controlling for childhood IQ scores. We focus on change in overall IQ given evidence that aging-related slopes are correlated across all cognitive functions, indicating that research on cognitive decline may be best focused on a highly reliable summary index, rather than focused on individual functions^94^.

#### Facial Age

Facial Age was based on two measurements of perceived age by an independent panel of eight people. First, age range was assessed by an independent panel of four raters, who were presented with standardized (non-smiling) digital facial photographs of Study members when they were 45 years old. Raters, who were kept blind to the actual age of Study members, used a Likert scale to categorize each Study member into a 5-year age range (i.e., from 20–24 years old up to 70+ years old). Interrater reliability was 0.77. Scores for each Study member were averaged across all raters. Second, relative age was assessed by a different panel of four raters, who were told that all photos were of people aged 45 years old. These raters then used a 7-item Likert scale to assign a “relative age” to each participant (i.e., 1 = “young looking,” to 7 = “old looking”). Interrater reliability was 0.79. The measure of perceived age at 45 years (i.e., Facial Age) was derived by standardizing and averaging age range and relative age scores.

### Human Connectome Project (HCP)

The HCP is a publicly available dataset that includes 1,206 participants with extensive MRI data^50^. HCP data access is managed by the WU-Minn HCP consortium. All participants provided informed consent. Specifically, we used data from 45 participants who completed the scan protocol a second time (with a mean interval between scans of approximately 140 days) allowing for calculation of test-retest reliability. All participants were free of current psychiatric or neurologic illness and were between 25 and 35 years of age. The mean age of the HCP test-retest sample analyzed here was 30.3 years (SD = 3.3 years, range = 22-35) at the first timepoint.

#### MRI

Structural MRI data were analyzed using the Human Connectome Project minimal preprocessing pipeline^85^. Briefly, T1-weighted images were processed using a custom FreeSurfer recon-all pipeline that is optimized for structural MRI with a higher resolution than 1 mm isotropic. Details on HCP MRI data acquisition have been described elsewhere^85^.

### Alzheimer’s Disease Neuroimaging Initiative (ADNI)

The primary goal of ADNI is to test whether serial MRI, PET, other biological markers, and clinical and neuropsychological assessments can be combined to measure the progression of neurodegeneration in participants with mild cognitive impairment, Alzheimer’s disease, and cognitively normal older adults (adni.loni.usc.edu)^95^. For further information, see adni.loni.usc.edu. Cognitive and diagnostic data were downloaded on June 12^th^, 2022. MRI data curated from the Alzheimer’s Disease Sequencing Project (ADSP) collection were downloaded on December 7^th^, 2023. ADNI was approved by the Institutional Review Boards of all the participating institutions. All participants provided written informed consent. ADNI sample demographic information can be found in **Supplemental Table S17.**

#### MRI

T1-weighted scans were collected using either 1.5T or 3T scanners. MRI acquisition parameters varied across ADNI sites and waves; however, the targets for acquisition were isotropic 1mm^3^ voxels.^96^. Raw T1-weighted images were processed using longitudinal FreeSurfer version 6.0. Scans were excluded for low quality if they did not have a QC rating of ‘Pass’ from ADNI investigators or if segmentation failed visual inspection. Scans were also excluded if participants were missing demographic data such as age, sex, or diagnosis (**Supplemental Figure S14**). Further details on MRI methods in ADNI can be found at adni.loni.usc.edu.

#### Cognitive and Behavioral Functioning

ADNI participants completed several cognitive and behavioral assessments at the time of scanning. The Alzheimer’s Disease Assessment Scale – Cognitive Subscale 13 (ADAS-Cog) is a structured scale that evaluates memory, reasoning, language, orientation, ideational praxis, and constructional praxis^97^. Delayed Word Recall and Number Cancellation are included in addition to the 11 standard ADAS Items^98^. The test is scored for errors, ranging from 0 (best performance) to 85 (worst performance). The Mini Mental State Exam (MMSE) is a screening instrument that evaluates orientation, memory, attention, concentration, naming, repetition, comprehension, and ability to create a sentence and to copy 2 overlapping pentagons^99^. The MMSE is scored as the number of correctly completed items ranging from 0 (worst performance) to 30 (best performance). The Montreal Cognitive Assessment (MoCA) is designed to detect people at the MCI stage of cognitive dysfunction^100^. The scale ranges from 0 (worst performance) to 30 (best performance). The Rey Auditory Verbal Learning Test is a list learning task which assesses learning and memory. On each of 5 learning trials, 15 unrelated nouns are presented orally at the rate of 1 word per second and immediate free recall of the words is elicited. After a 30-minute delay filled with unrelated testing, free recall of the original 15-word list is elicited. Both immediate recall and the percent forgotten are used. The Logical Memory tests I and II (Delayed Paragraph Recall) is from the Wechsler Memory Scale–Revised. Free recall of 1 short story is elicited immediately after being read aloud to the participant and again after a 30-minute delay. The total bits of information recalled after the delay interval (maximum score = 25) are analyzed. The Trail Making Test, Part B, consists of 25 circles, either numbered (1 through 13) or containing letters (A through L). Participants connect the circles while alternating between numbers and letters (e.g., A to 1; 1 to B; B to 2; 2 to C). Time to complete (300 seconds maximum) is the primary measure of interest. The Functional Assessment Questionnaire (FAQ) is a self-report measure of instrumental activities of daily living such as preparing meals, performing chores, keeping a schedule, and traveling outside of one’s neighborhood^101^. Each unique cognitive testing measure was paired with the participant’s most temporally proximate brain scan within 6 months of cognitive testing.

#### Cognitive Status

ADNI participants were classified into cognitively normal (CN), mild cognitive impairment (MCI), or dementia groups by ADNI study physicians based on subjective memory complaints, multiple neurocognitive and behavioral assessment scores, and level of impairment in activities of daily living. Complete diagnostic criteria can be found at adni.loni.usc.edu. Each individual scan was categorized according to the most temporally proximate cognitive diagnosis received by that participant.

#### Education

Education level was measured according to self-reported years of education. For the purposes of visualization in **Figure 5E**, participants were grouped according to the following thresholds: Less than high school: < 12 years; High school: 12 years; Some college: 12-15 years; College: 16 years; More than college: >16 years.

### UK Biobank

The UK Biobank is a United Kingdom population-based prospective study of 502,486 participants between the ages of 40 and 69 at baseline assessment^102^. We analyzed data from 42,583 participants who underwent brain MRI. Data used in these analyses were downloaded in April 2023. The UK Biobank was approved by the North West Multi-centre for Research Ethics Committee. All participants provided written informed consent. UK Biobank sample demographic information can be found in **Supplemental Table S17**.

#### MRI

MRI methods for the UK Biobank have been described in detail elsewhere^103^. Briefly, MRI data were collected using 3 identical 3T Siemens Skyra scanners with a 32-channel Siemens head coil. T1-weighted images were obtained using a 3D MP-RAGE with the following parameters: TR = 2000 ms; TI = 880 ms; 208 sagittal slices, matrix =256×256; slice thickness = 1 mm with no gap; and total scan time = 4 min and 52 s. Our study made use of imaging-derived phenotypes generated by an image-processing pipeline developed and run on behalf of the UK Biobank^103^. As part of this pipeline, raw T1-weighted images were processed using the cross-sectional FreeSurfer version 6.0. All brain measures used in the cross-sectional analyses presented here were derived from the outputs of this FreeSurfer pipeline. We excluded UK Biobank participants with very low signal-to-noise ratio and highly unusual summary morphometrics indicative of low-quality reconstruction **(Supplemental Figure S15**).

To measure hippocampal volume change among the subset of UK Biobank participants with longitudinal MRI data, we reprocessed all T1-weighted images for this subset of participants using the longitudinal FreeSurfer version 6.0 pipeline^104^. This allowed us to avoid the known biases that can be introduced by different processing stages of the longitudinal pipeline on different hardware and software environments. Specifically, we reprocessed both time points of each participant’s T1-weighted scans with the cross-sectional recon-all pipeline^105^. Then we built an unbiased within-subject template^106^ using robust, inverse consistent registration^107^ and reprocessed each T1-weighted scan through the automated longitudinal pipeline^104^.

#### Cognitive Functioning

UK Biobank participants completed a battery of cognitive tests at the time of MRI. We investigated cognitive functioning using the following measures: Reaction Time (Field ID = 20023), Fluid Intelligence (Field ID = 20016), Numeric Memory (Field ID = 4282), Trails A (Field ID = 6348) and B (Field ID = 6350), Symbol Digit Substitution (Field ID = 23324), Tower Rearranging (Field ID = 21004), and Matrix Completion (Field ID = 6373). The details of these cognitive tests have been described elsewhere^108^.

#### Frailty and Self-Reported Health

To further investigate aging-related health, we used the Fried Frailty Index^54^. Briefly, the Fried Frailty Index is based on meeting criteria for declining functioning across five domains: unintentional weight loss, exhaustion, weakness, physical inactivity, and slow walking speed. Index scores range from 0 to 5 with higher scores indicating greater frailty^109^. During their imaging visit, UK Biobank participants were also asked to rate their overall health as “Poor,” “Fair,” “Good,” or “Excellent.” We used these ratings to investigate self-reported overall health (Field ID = 4548).

#### Disease and Mortality Records

To assess the influence of DunedinPACNI on aging-related disease and mortality risk in UK Biobank we used variables from algorithmically defined health outcomes. Briefly, algorithmically defined outcomes are generated by combining information from baseline assessments (self-reported medical conditions, operations, and medications) with linked data from hospital admissions and death registries. Due to the relatively small number of aging-related disease diagnoses at follow-up, we defined aging-related morbidity as being diagnosed with myocardial infarction (Field ID = 42000), chronic obstructive pulmonary disease (Field ID = 42016), dementia (Field ID = 42018), or stroke (Field ID = 42006). Furthermore, we defined risk of chronic disease as the emergence or one or more of these diagnoses among participants who were healthy at the time of scanning (i.e., baseline). Mortality was quantified during follow-up from death records (Field ID = 40000).

#### Education, Income, and Ethnicity

To test the association between DunedinPACNI and socioeconomic gradients of health, we tested whether UK Biobank participants differed in DunedinPACNI scores as a function of educational attainment and household income. We grouped participants into three categories according to their self-reported educational qualifications (Field ID = 6138) following prior work^110^. Specifically, these groups were: high (college or university degree), medium (A/AS levels or equivalent or O/levels/GCSEs or equivalent), and low (none of these). We also tested whether UK Biobank participants differed in DunedinPACNI scores as a function of household income (Field ID = 738).

We also conducted sensitivity analyses while restricting the UK Biobank sample to either only low income or only non-White participants. We considered participants low income if they reported making < £18,000 per year in household income. We considered participants to be non-White if they did not report their ethnic background (Field ID = 21000) as “Any other white background,” “British,” “Do not know,” “Irish,” “Prefer not to answer,” or “White.”

### Latin American Brain Health Institute (BrainLat)

BrainLat is a multimodal neuroimaging dataset of patients with neurodegenerative diseases and healthy adult controls collected in Argentina, Chile, Colombia, Mexico and Peru^64^. We analyzed neuroimaging data from 368 individuals who were either cognitively healthy or diagnosed with Alzheimer’s disease or behavior variant frontotemporal dementia. The BrainLat Study was approved by the institutional ethical boards of each recruitment site. All participants, or their legal representatives, provided written informed consent. BrainLat demographic data can be found in **Supplemental Table S17.**

#### MRI

MRI methods for BrainLat have been described in detail elsewhere^64^. Briefly, T1-weighted MPRAGE scans were collected on either 1.5 or 3T scanners. Acquisition parameters varied across sites, but scans most frequently had isometric 1mm^3^ voxels. Scans were downloaded and then processed using FreeSurfer version 6.0. Participants were excluded if they failed automated FreeSurfer quality metrics or a visual QC of segmentation output (**Supplementary Figure S16**).

#### Diagnostic Classification

All included participants could speak fluent Spanish and had adequate visual and auditory capacity for testing. Participants were classified as cognitively normal if they had a modified clinical dementia rating of 0, an MMSE score above 25, and lacked history of substance abuse, neurological, or psychiatric disorders. Patients were classified into the Alzheimer’s disease (AD) or frontotemporal dementia (FTD) groups according to the National Institute of Neurological Disorders and Stroke – Alzheimer Disease and Related Disorders working group for probable Alzheimer’s disease or probable behavioral variant frontotemporal dementia. Diagnosis was supported using appropriate MRI or PET imaging when needed^64^.

#### Cognitive Status

BrainLat participants were evaluated with the Montreal Cognitive Assessment (MoCA). The MoCA is designed to detect people at the MCI stage of cognitive dysfunction^100^. The scale ranges from 0 (worst performance) to 30 (best performance).

## STATISTICAL ANALYSES

### Pace of Aging

The derivation of the Pace of Aging has been described elsewhere^1,2^. Briefly, we measured a panel of the following 19 biomarkers (**Figure 1A**) at ages 26, 32, 38, and 45: body mass index (BMI), waist-hip ratio, glycated hemoglobin, leptin, blood pressure (mean arterial pressure), cardiorespiratory fitness (VO_2_max), forced vital capacity ratio (FEV_1_/FVC), forced expiratory volume in one second (FEV_1_), total cholesterol, triglycerides, high-density lipoprotein (HDL), lipoprotein(a), apolipoprotein B100/A1 ratio, estimated glomerular filtration rate (eGFR), blood urea nitrogen (BUN), high sensitivity C-reactive protein (hs-CRP), white blood cell count, mean periodontal attachment loss (AL), and the number of dental-caries-affected tooth surfaces (tooth decay). To calculate each Study members Pace of Aging, we first transformed the biomarker values to a standardized scale. For each biomarker at each wave, we standardized values according to the age-26 distribution. Next, we calculated each Study member’s slope for each of the 19 biomarkers using a mixed-effects growth model that regressed the biomarker’s level on age. Finally, we combined information from the 19 slopes of the biomarkers using a unit-weighting scheme. We calculated each Study member’s Pace of Aging as the sum of age-dependent annual changes in biomarker Z-scores. Biomarker standardization was performed separately for men and women.

### DunedinPACNI

A schematic of DunedinPACNI model development can be found in **Figure 1**. We trained an elastic net regression model to estimate the Pace of Aging from structural neuroimaging phenotypes in 860 Dunedin Study members at age 45 (for attrition analysis and inclusion criteria see **Supplemental Figures S1-S2, S12**). We selected 315 FreeSurfer measures as predictors from the following categories: regional cortical thickness (CT), regional cortical surface area (SA), regional cortical gray matter volume (GMV), regional cortical gray-white matter signal intensity ratio (GWR), and ‘ASEG’ volumes (i.e., regional subcortical gray matter volumes, ventricular volumes, and bilateral volume of white matter hypointensities). All cortical data were parcellated according to the Desikan-Killiany Atlas^111^. Note that although there are many ADNI scans that do not pass QC (**Supplemental Figure S14**), FreeSurfer is a robust segmentation method, especially among healthy individuals ^112^. Four phenotypes from the ‘ASEG’ volumes were excluded due to insufficient variance in the Dunedin Study (left and right white matter hypointensities, left and right non-white matter hypointensities). Model training was performed using the *caret* package in R. We conducted a grid search across a range of α and α; values. We used 100 repetitions of 10-fold cross-validation to estimate model performance in held-out participants. The effect of sex was regressed from the Pace of Aging prior to model training. To prevent information leak during cross-validation, we regressed sex from each training set and applied the resulting beta weights to each test set. This approach ensured that our model only used information from the training set, including covariate regression, when calculating predictions in each test set. We selected optimal tuning parameters according to highest variance explained and lowest mean absolute error. The optimal tuning parameters were α = 0.214 and λ = 0.100. Using these parameters, we fit the model to the entire N=860 sample. The raw elastic net regression model weights can be found in **Supplemental Table S18**.

To generate DunedinPACNI scores in HCP, ADNI, and UK Biobank participants, we applied the regression weights from the DunedinPACNI model to FreeSurfer-derived phenotypes within each dataset and summed the products and model intercept. In ADNI and UK Biobank, DunedinPACNI scores were correlated with chronological age (ADNI: r=0.37; UK Biobank: r=0.50; BrainLat: r=0.37; **Supplemental Figure S17**).

In addition, we conducted the same procedure again without GWR as this measure is not always distributed in public datasets. We observed slightly reduced model accuracy when GWR was not included. DunedinPACNI estimates without GWR phenotypes showed excellent test re-test reliability in HCP. DunedinPACNI estimates were similar with and without GWR phenotypes in ADNI, UK Biobank, and BrainLat (see **Supplemental Figure S18** for more details).

### Brain Age Gap

We submitted raw T1-weighted images from ADNI, UK Biobank, and BrainLat to the publicly available brainageR algorithm. This model, which has been described in detail elsewhere^113^, is trained to predict chronological age among a sample of healthy, cognitively unimpaired individuals aged 18-92 years. This algorithm was selected because it generates the most reliable estimates amongst published algorithms^65^. Briefly, brainageR is estimated by first segmenting and normalizing T1-weighted images using SPM12. Next, coefficients derived from a Gaussian Process regression model predicting chronological age in a training dataset (N=2,001) are applied to morphometric features from brain segmentations to predict participants’ chronological age. Brain age gap was subsequently estimated by subtracting actual chronological age from predicted age^113^. Of note, 15 ADNI scans failed the brain age gap pipeline (14 failed visual inspection of segmentation, 1 error computing predicted age). These scans were excluded from all brain age gap analyses, including comparative analyses with DunedinPACNI.

### Dunedin Study Validation Analyses

To first test the validity of DunedinPACNI within the Dunedin Study training sample, we tested for linear associations between DunedinPACNI scores and one-legged balance, gait speed, step in place, chair stands, grip strength, visual-motor coordination, subjective physical limitations, subjective health, cognitive function, child-to-adult cognitive decline, and facial aging while controlling for sex. We compared these effect sizes to associations between each of these measures and the original, 20-year Pace of Aging.

### Test-Retest Reliability

We used the HCP dataset to assess the test-retest reliability of DunedinPACNI. Reliability was quantified using a two-way mixed-effects ICC (3,1) with session modeled as a fixed effect, subject as a random effect, and test-retest interval as an effect of no interest^114^.

### Cognitive and Physical Functioning

We first used linear regression models to test for associations between DunedinPACNI and scores on tests of cognition, physical function, and health in ADNI and UK Biobank. All analyses controlled for age and sex. In ADNI, we calculated robust standard errors to account for non-independence from repeated observations. We also tested the standardized differences in DunedinPACNI scores between three groups based on cognitive status: cognitively normal (CN), mild cognitive impairment (MCI), and dementia. All group difference comparisons controlled for age and sex. We again calculated robust standard errors to account for non-independence and conduced a sensitivity analysis while controlling for *APOE* ε4 carriership. We repeated these analyses with brain age gap. Of note, when conducting analyses on the combined effects of DunedinPACNI and brain age gap on cognitive outcomes in ADNI, we restricted the sample to the first timepoint of each measure. This included only one observation per participant, allowing us to more easily combine effect sizes and confidence intervals.

### Dementia Survival Analysis

We conducted a Cox-proportional hazard regression using ADNI participants’ baseline DunedinPACNI scores to predict their probability of cognitive decline or clinically conversion to dementia during the follow-up window. Conversion among cognitively normal participants was defined as having a diagnosis of CN at baseline but a diagnosis of MCI or dementia at the end of follow-up. Conversion among MCI participants was defined as having a diagnosis of MCI at baseline and a diagnosis of dementia by the end of follow-up. Participants who had a baseline diagnosis of dementia or transitioned from MCI to CN were not included in this analysis. The analysis controlled for sex, age at baseline, and length of observation window. We investigated the influence of AD genetic risk on these results by conducting all analyses while additionally controlling for *APOE* ε4 carriership. We repeated these analyses for brain age gap.

### Prediction of Hippocampal Atrophy Rates

We used repeated MRI measurements from the ADNI (N = 1,302) and UK Biobank (N = 4,628) to generate estimates of change in hippocampal gray matter volume. We ran longitudinal ComBat on ADNI MRI data to remove differential scanner effects^115^. Next, using all available timepoints for each participant, we generated multilevel linear models for bilateral hippocampal volume with random effects for both participant and age. Using these models, we derived trajectories to track change in hippocampal gray matter volume for each participant. We then tested whether each participant’s baseline DunedinPACNI scores could predict their subsequent rate of hippocampal atrophy. These analyses controlled for age, sex, and length of observation period. We investigated the influence of AD genetic risk on these results by conducting these analyses while additionally controlling for *APOE* ε4 carriership. We repeated these analyses for brain age gap.

### Morbidity and Mortality Survival Analyses

To investigate the association between DunedinPACNI and morbidity, we used UK Biobank data to calculate the standardized differences in DunedinPACNI scores between three groups based on number of lifetime chronic disease diagnosis (0, 1, 2+). Next, we conducted a Cox-proportional hazard regression using UK Biobank participants’ baseline DunedinPACNI scores to predict the onset of a chronic aging-related disease (N=827 emergent diagnoses: myocardial infarction, chronic obstructive pulmonary disease, dementia, or stroke) in participants who had never previously received any of these diagnoses at the time of scanning (N=40,753). Similarly, to investigate the association between DunedinPACNI and mortality we conducted a Cox-proportional hazard regression using UK Biobank participants’ baseline DunedinPACNI scores to predict death (N=757 deaths). Both models controlled for baseline age, time to onset, and sex. We repeated these analyses for brain age gap.

### Socioeconomic Inequality Analyses

To investigate whether DunedinPACNI reflected gradients of socioeconomic inequality^58^, we first tested for linear relationships between DunedinPACNI and years of education in ADNI and UK Biobank. We also tested for a linear relationship between DunedinPACNI and household income in UK Biobank. These analyses controlled for sex and age. In ADNI, we included only the first MRI observation per participant.

### Replication in a Latin American sample

To investigate whether DunedinPACNI generalizes to samples of individuals who are underrepresented in neuroimaging research^60^, we tested whether the degree of acceleration in DunedinPACNI among ADNI participants with dementia was similar in BrainLat participants with dementia. We tested for standardized differences by comparing the AD and FTD groups to the CN group, respectively. We also tested for linear associations between DunedinPACNI and MoCA scores. All analyses in this sample controlled for age and sex. We then compared the magnitude of acceleration among BrainLat dementia patients to the previously identified acceleration among ADNI dementia patients. Lastly, we compared the strength of the linear association between DunedinPACNI and MoCA scores among BrainLat participants to the previously identified association in ADNI participants.

### Comparison with hippocampal and ventricular volume

We investigated how DunedinPACNI differs from two commonly used MRI-based measures of brain aging: hippocampal volume and ventricular volume. We calculated hippocampal volume as the sum of left and right hippocampus volume measures derived from FreeSurfer. Likewise, we calculated ventricular volume as the sum of the left and right lateral ventricle volume measures. We first repeated cross-sectional associations with cognition, frailty, and poor health in the UK Biobank substituting DunedinPACNI with hippocampal volume or ventricular volume. Next, we conducted this same procedure with our Cox-proportional hazard regression models of chronic disease and mortality risk in UK Biobank, and cognitive decline risk among cognitively normal ADNI participants. Lastly, we compared coefficients for all analyses while including DunedinPACNI and either hippocampal volume or ventricular volume in respective models. All analyses controlled for age and sex.

## DATA AVAILABILITY

Dunedin Study data is available via managed access at https://sites.duke.edu/moffittcaspiprojects/data-use-guidelines/. The Human Connectome Project data are publicly available at http://www.humanconnectomeproject.org/data/. Alzheimer’s Disease Neuroimaging Initiative data are publicly available at https://adni.loni.usc.edu/. Researchers can apply to access all UK Biobank data at https://ams.ukbiobank.ac.uk/ams/. BrainLat data is publicly available at https://www.synapse.org/Synapse:syn51549340/wiki/624187.

## CODE AVAILABILITY

The DunedinPACNI algorithm will be made available for download upon publication. All scripts used in the analyses presented here are available at https://github.com/etw11/WhitmanElliott_2024.

## ACKNOWLEDGEMENTS

This research received support from the US National Institute on Aging (grants R01AG049789, R01AG032282, R01AG073207) and the UK Medical Research Council (grant MR/X021149/1). The Dunedin Multidisciplinary Health and Development Research Unit is supported by the New Zealand Health Research Council (Programme Grant 16-604).

We thank the Dunedin Study members, Unit research staff, previous Study Director, Emeritus Distinguished Professor, the late Richie Poulton, for his leadership during the Study’s research transition from young adulthood to aging (2000-2023), and Study founder Dr Phil A. Silva. The Dunedin Unit is located within the Ngāi Tahu tribal area who we acknowledge as first peoples, tangata whenua (people of this land).

This research has been conducted using the UK Biobank Resource under Application Number 67237.

Data collection and sharing for the Alzheimer’s Disease Neuroimaging Initiative (ADNI) is funded by the National Institute on Aging (National Institutes of Health Grant U19 AG024904). The grantee organization is the Northern California Institute for Research and Education.

In the past, ADNI has also received funding from the National Institute of Biomedical Imaging and Bioengineering, the Canadian Institutes of Health Research, and private sector contributions through the Foundation for the National Institutes of Health (FNIH) including generous contributions from the following: AbbVie, Alzheimer’s Association; Alzheimer’s Drug Discovery Foundation; Araclon Biotech; BioClinica, Inc.; Biogen; Bristol-Myers Squibb Company; CereSpir, Inc.; Cogstate; Eisai Inc.; Elan Pharmaceuticals, Inc.; Eli Lilly and Company; EuroImmun; F. Hoffmann-La Roche Ltd and its affiliated company Genentech, Inc.; Fujirebio; GE Healthcare; IXICO Ltd.; Janssen Alzheimer Immunotherapy Research & Development, LLC.; Johnson & Johnson Pharmaceutical Research &Development LLC.; Lumosity; Lundbeck; Merck & Co., Inc.; Meso Scale Diagnostics, LLC.; NeuroRx Research; Neurotrack Technologies; Novartis Pharmaceuticals Corporation; Pfizer Inc.; Piramal Imaging; Servier; Takeda Pharmaceutical Company; and Transition Therapeutics.

We thank the Latin American Brain Health Institute and associated investigators for sharing the BrainLat dataset.

## AUTHOR CONTRIBUTIONS

E.T.W., M.L.E., A.R.K., A.C., T.E.M, and A.R.H. designed the research. E.T.W., M.L.E., A.R.K., W.C.A., T.J.A., N.C., S.H., D.I., T.R.M., S.R., K.S., R.T., B.S.W., A.C., T.E.M., and A.R.H performed the research. E.T.W., M.L.E., and A.R.K. analyzed data. E.T.W., M.L.E., A.R.K., A.C., T.E.M., and A.R.H. wrote the paper.

